# Chemogenomic screening Identifies the Hsp70 Co-chaperone HDJ2 as a Hub for Anticancer Drug Resistance

**DOI:** 10.1101/818427

**Authors:** Nitika, Jacob S. Blackman, Laura E. Knighton, Jade E. Takakuwa, Stuart K. Calderwood, Andrew W. Truman

## Abstract

Heat shock protein 70 (Hsp70) is an important molecular chaperone that regulates oncoprotein stability and tumorigenesis. However, attempts to develop anti-chaperone drugs targeting molecules such as Hsp70 have been hampered by toxicity issues. Hsp70 is regulated by a suite of co-chaperone molecules that bring “clients” to the primary chaperone for efficient folding. Therefore, rather than targeting Hsp70 itself, here we have examined the feasibility of inhibiting the co-chaperone HDJ2, a member of the J domain protein family, as a novel anticancer strategy. We found HDJ2 to be upregulated in a variety of cancers, suggesting a role in malignancy. To confirm this role, we screened the NIH Approved Oncology collection for chemical-genetic interactions with loss of HDJ2 in cancer. 41 compounds showed strong synergy with HDJ2 loss, whereas 18 dramatically lost potency. Several of these hits were validated using a HDJ2 inhibitor (116-9e) in castration-resistant prostate cancer cell (CRPC) and spheroid models. Taken together, these results confirmed that HDJ2 is a hub for anticancer drug resistance and that HDJ2 inhibition may be a potent strategy to sensitize cancer cells to current and future therapeutics.

**Graphical Abstract:** HDJ2 knockout or inhibition via small molecule impacts cellular resistance to anticancer therapeutics.

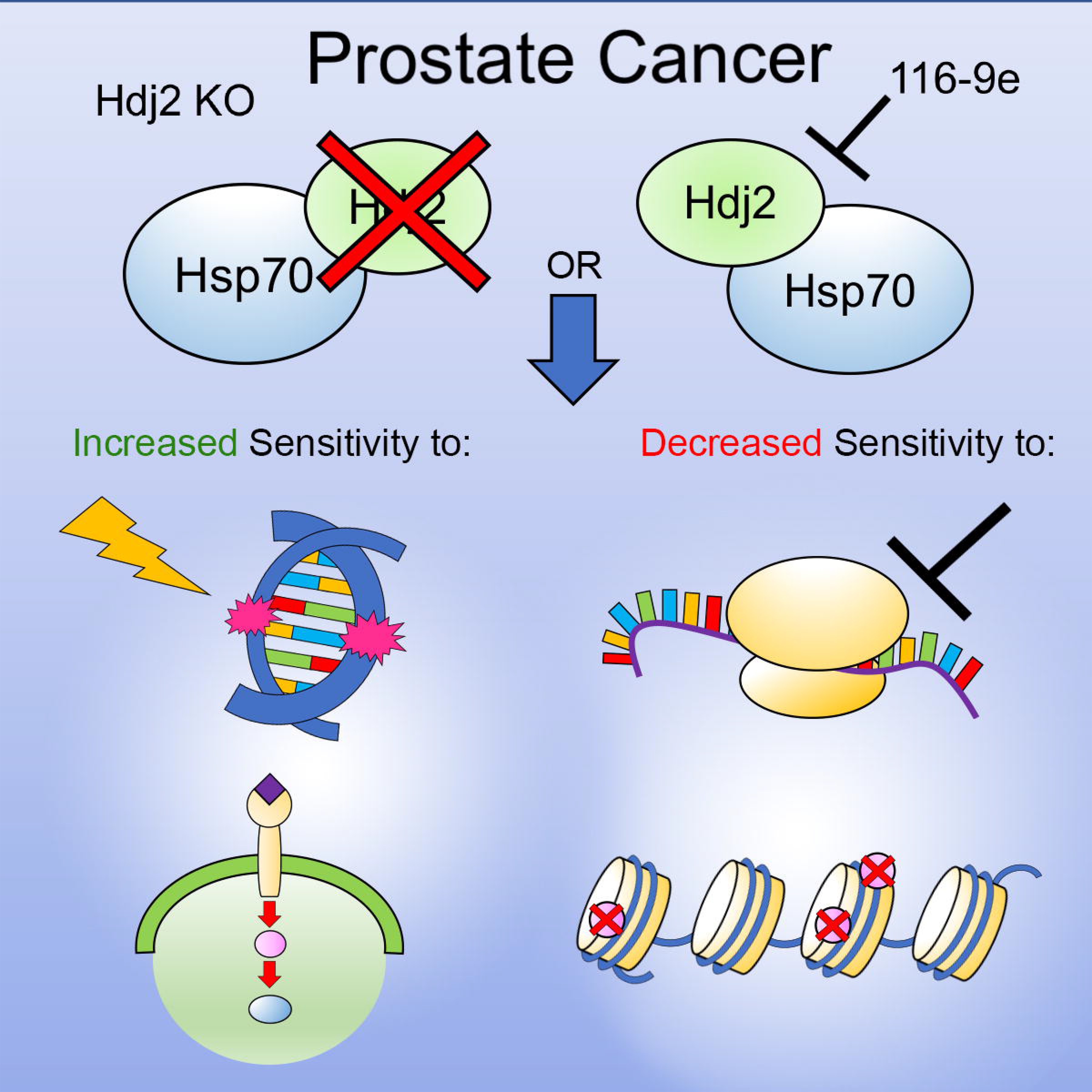

## Introduction

Hsp70 is a molecular chaperone that plays important roles in protein quality control processes such as protein folding, transport, degradation, regulation and aggregation prevention [1]. Hsp70 levels are elevated in various cancers and overexpression correlates with poor prognosis for survival and response to cancer therapy [2]. The elevated levels of Hsp90 and Hsp70 chaperones in cancer and their role in fostering multiple oncogenic pathways has made these proteins attractive drug targets with numerous anti-chaperone compounds having been developed so far [3]. Problematically, Hsp70 is required for cell survival and protein homeostasis, and thus its inhibition is detrimental to the viability of both normal and cancer cells, with dubious selectivity for tumor cells [4].

Hsp70 performs all its functions in association with a large spectrum of helper proteins known as co-chaperones that include J-proteins, tetratricopeptide repeat (TPR) domain-containing proteins and nucleotide exchange factors (NEFs) which fine-tune Hsp70 specificity and activity in the cell. The J-proteins recruit the protein substrates or clients and interact with such clients at the interface of NBD and SBDβ of Hsp70 [5]. This interaction leads to increased Hsp70-mediated ATP turnover and activation of protein folding. J-proteins have a highly conserved 70 amino acid motif containing Histidine, Proline and Aspartic acid amino acid residues known as HPD motif which is essential for stimulating ATPase activity of Hsp70 [6]. In humans, the J-protein family has about 50 members which are further divided into three groups based on the localization of J domain within a protein [7, 8]. DNAJA1 (more commonly referred to as HDJ2) associates with unfolded polypeptide chains, preventing their aggregation [7]. Several Hsp70 inhibitors have failed in clinical trials due to their toxicity. More recently, alternative strategies have focused on sensitizing cells to anticancer agents by either manipulating post-translational modification of chaperones or their interaction with specific co-chaperones [4, 9–19]. HDJ2 (mammalian homologue of yeast Ydj1) is an interesting possible anticancer target as a key mediator of Hsp70 function that appears to regulate specific features of tumorigenesis [13, 20, 21]. A recent study demonstrated that CRPCs expressing ARv7 are insensitive to Hsp90 inhibitors but are sensitive to Hsp40 inhibition [22]. In addition, we have shown that targeting specific oncoprotein complexes (ribonucleotide reductase) with a combination of traditional as well as an HDJ2 inhibitor produces highly synergistic effects [13]. We propose that targeting HDJ2 in cancer may offer an attractive alternative to the toxicity induced by full Hsp90/Hsp70 inhibition.

Anticancer monotherapies using broadly active cytotoxic or molecularly targeted drugs are limited in their ability to demonstrate a reliable clinical response. This is due to redundant signaling pathways, feedback loops and resistance mechanisms in the cancer cells [23, 24]. Thus, combination anticancer therapies have been used clinically for over 50 years to improve the responses achieved by monotherapies alone. Cancer cell line-based models for these combination therapies are easy and inexpensive to perform using high-throughput drug screening protocols (HTS) to identify the most effective drug combination [25, 26]. HTS helps to explore the relation between the cell line characteristics and drug specific dose responses [25]. Chemogenomics is one such HTS based approach where large collection of anticancer chemical drugs are screened to identify biological targets. These screening sets often contain small molecules that are well annotated and have defined molecular targets. Such an approach is particularly beneficial for cancer research because malignant cells often contain multiple aberrations which require targeted therapy to inactivate cancer driver activities and mitigate deleterious effects of the drugs to normal cells [24, 27].

Here, we performed an unbiased screen of the NIH Approved Oncology Drug set containing 131 anti-cancer drugs in combination with HAP1 cancer cell lines depleted of J-protein HDJ2. We identified 41 compounds showing strong synergy with the loss of HDJ2, and by contrast 18 molecules displaying reduced potency in the knockout cell line. We validated three drugs (cabozantinib, clofarabine and vinblastine) in combination with a unique HDJ2 inhibitor (116-9e) for synergy in the LNCaP cancer cell lines and confirmed omacetaxine mepesuccinate, idarubicin and sorafenib for antagonism (i.e. with reduced potency after HDJ2 inhibition). This study demonstrates the validity of developing Hsp70 co-chaperone inhibitors to sensitize cells to current anticancer therapies and suggests that determining HDJ2 status of a tumor may be beneficial in selecting the most appropriate course of treatment.

## Materials and Methods

### Cell culture

The HAP1 Chronic Myelogenous Leukemia cancer cell line and HDJ2 Knockout cell line was purchased from Horizon Discovery and were cultured in Iscove’s Modified Eagle Medium (Invitrogen) with 10% fetal bovine serum (Gibco), 100 units/ml penicillin, and 100 μg/ml streptomycin at 5% CO_2_ and 37° C. The LNCaP cancer cell line was purchased from ATCC and were cultured in RPMI-1640 medium (Invitrogen) with 10% fetal bovine serum (FBS, Clontech), 100 units/ml penicillin, and 100 μg/ml streptomycin at 5% CO_2_ and 37° C.

### Drug Screening

Approved Oncology Drug plates consisting of the most current FDA approved anticancer drugs were obtained from National Cancer Institute (NCI). For experiments delineating the synergy between the loss of HDJ2 and approved anticancer drug, HAP1 cells and HAP1 (HDJ2 KO) cells were plated in growth media at 20% confluency 1 day prior to drug treatment. On Day 1 of treatment, cells were treated with DMSO (control), Approved oncology anticancer drugs at 50 μM for 72 hours. Following drug treatments, Cell Titer-Glo reagent was added directly to the wells according to manufacturer’s instructions. The luminescence was measured on Bio-Tek Plate reader. Luminescence reading was normalized to and expressed as a relative percentage of the plate averaged DMSO control. The data shown are the mean and SEM of three independent biological replicates.

### Combination index (CI) calculations

For IC_50_ calculations, LNCaP cells were seeded in triplicates in 96-well white bottom Nunc plates in growth media at 20% confluency 1 day prior to initiation of drug treatment. On Day 1 of treatment, cells were treated with DMSO (control) and ten folds serial dilution of anti-cancer drugs Cabozantinib, Clofarabine, Vinblastine, Sorafenib, Idarubicin and Omacetaxine mepesuccinate and 116-9e. After 72 h, cell viability was measured using Promega Cell Titer-Glo cell viability assay on Bio-Tek plate reader. The combination index was calculated using the Chou-Talalay method using CompuSyn software[28].

### Spheroid Generation

Single-cell suspensions (5000/well) were plated in one well of 24-well plates in a 1:1 mixture of RPMI medium and Matrigel (BD Bioscience CB-40324). Cells in Matrigel are kept cold at all times and under continuous agitation. Warm PBS is added to all empty wells, if any. Plates are incubated at 37 °C with 5% CO_2_ for 15 min to solidify the gel before addition of 100 μl of pre-warmed RPMI to each well. Two days after seeding, medium is fully removed and replaced with fresh RPMI containing the indicated drugs. The same procedure is repeated daily on two consecutive days. Twenty-four hours after the last treatments, media is removed and wells are washed with 100 μl of pre-warmed PBS. To prepare for downstream assays, spheroids are then released from Matrigel by incubating at 37 °C for 40 min in 100 μl of 10 mg/mL dispase (Sigma).

### Apoptosis assay

Apoptosis of LNCaP spheroids was detected by the Annexin V–FITC/propidium iodide–binding assay. Cells were treated with either 0.1% DMSO (dimethyl sulfoxide),116-9e, Cabozantinib, Clofarabine, Vinblastine, Sorafenib, Idarubicin, Omacetaxine mepesuccinate and Sorafenib alone or in combination with 116-9e for 48 hours at the IC_50_ concentrations, and then stained with Annexin V–FITC and propidium iodide. The rate of apoptosis was determined using BD FORTESSA, and data were analyzed using FlowJo software and were reported as the mean ± SD. The results are representative of three independent experiments.

### Bioinformatics

Cancer genome data and Cancer Cell Line Encyclopedia data were accessed from the cBioPortal (www.cbioportal.org) for Cancer Genomics (Gao et al, 2013). Total patient numbers and detailed information regarding published datasets and associated publications are indicated in Fig 1A and 1B.

**Figure 1.**
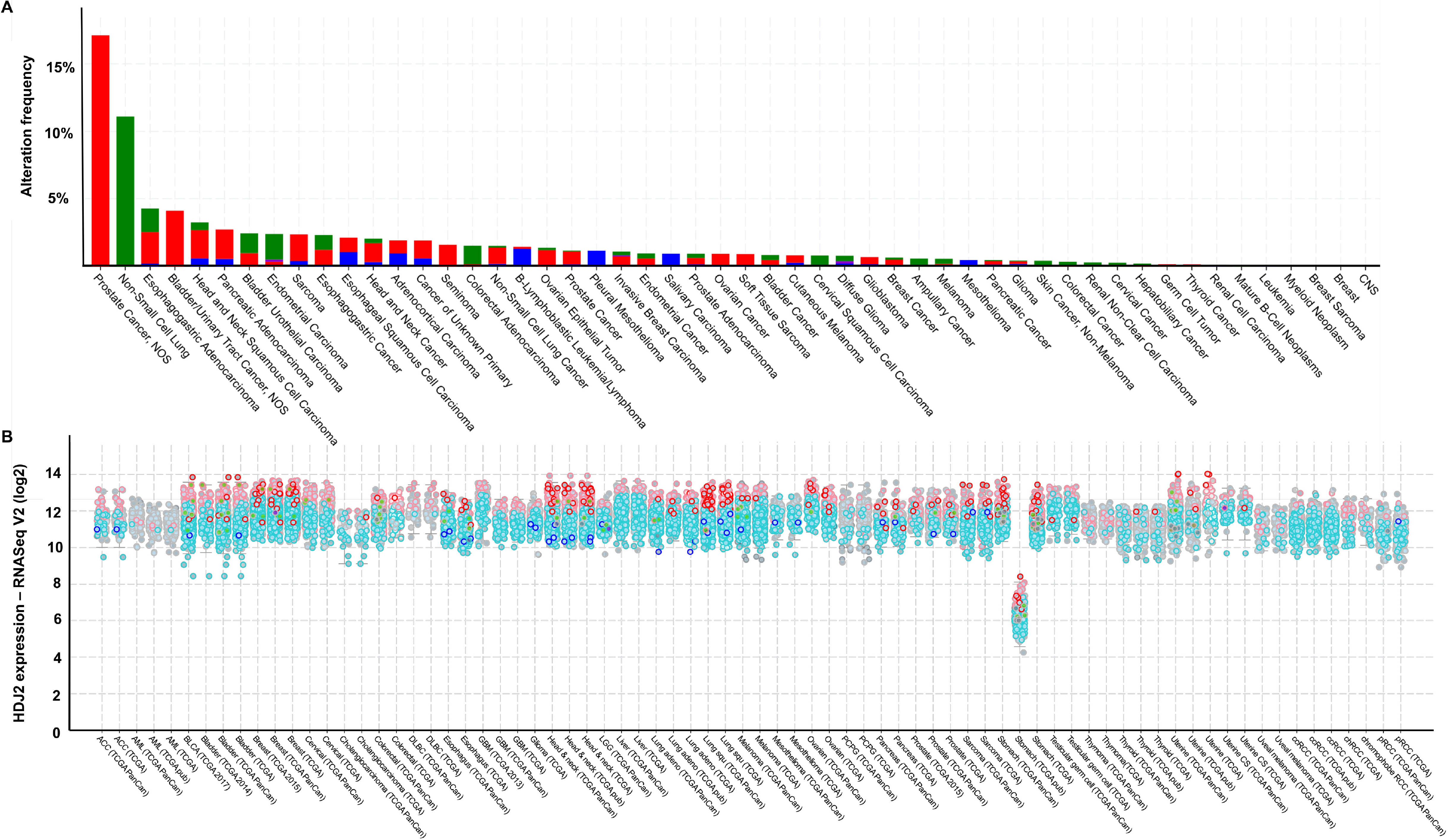
HDJ2 is altered in cancer. (A) Prevalence of HDJ2 alterations in various cancer genomes analyzed via the cBioPortal. Red bar, amplification. Blue bar, homozygous deletion. Green square, missense mutation. Purple square, Fusion. (B) HDJ2 mRNA expression in tumor determined via cBioPortal. P-value for a gene represents its P-value for the median-ranked analysis.

### Statistical analysis

Data were analyzed using GraphPad Prism built-in statistical tests indicated in relevant figure legends. The following asterisk system for P value was used: P <0.05; P <0.01; 0.001; and P <0.0001.

## Results

### HDJ2 is mutated and overexpressed in a variety of cancers

We first investigated the incidence of HDJ2 alterations in cancer using cancer genomics databases. Mutations and copy number changes occur in HDJ2 at a relatively low level (<5% of samples) in the majority of cancer types (Figure 1). However, the data shows that HDJ2 is strikingly amplified in neuroendocrine prostate cancer (NEPC) at a frequency of 18.42% (Figure 1A). Additionally, HDJ2 is mutated in 11.1% of Non-Small Cell Lung Cancer (NSCLC) cases (Figure 1A). Hsp70 and Hsp90 are often overexpressed in tumors [2, 29, 30]. To determine if the HDJ2 gene is also overexpressed in cancer, we analyzed the expression data from 72,175 samples in 236 studies (cBioportal database) [31, 32]. Interestingly, HDJ2 was expressed at significantly higher levels in cancer samples, with a median expression in cancer of between log2 values of 10 and 14 (Figure 1B). Taken together, these data suggest that alteration of HDJ2 function may be important in the malignant properties of cancer cells.

### Characterizing the role of HDJ2 in anticancer drug resistance

The existing literature is contradictory as to whether HDJ2 may possess tumor suppressor or driver properties [21, 33].To clarify whether silencing of HDJ2 could be beneficial in the treatment of cancer, we screened wildtype HAP1 cells and HAP1 cells lacking HDJ2 (HAP1^HDJ2 KO^) for comparative resistance against the NIH NCI Approved Oncology Collection (Figure 2A) (https://dtp.cancer.gov/organization/dscb/obtaining/available_plates.html). According to pharmacologic action, the compounds in the library have been divided into seven categories: Protein synthesis inhibitors, Proteasome inhibitors, Epigenetic modifiers, Metabolic inhibitors, Cytoskeletal inhibitors, Signal transduction inhibitors and DNA synthesis and repair inhibitors. Further fold enrichment of each drug category was calculated for the drugs whose potency increased or decreased with HDJ2 KO. To monitor the screening quality, each screening plate contained control wells treated with vehicle (1% DMSO). The final concentration of the screening compounds was 50 μmol/L. Positive hits (synergistic) or negative hits (antagonistic) were determined by normalizing the log_2_ ratio of viability of HDJ2 knockout cells over wildtype cells. A full list of the screening results is shown in Supplementary Table T1 and the sorted data are graphically plotted in Figure 2B. 41 drugs had increased potency upon HDJ2 deletion whereas 18 drugs displayed reduced potency. Drug target analysis was carried out by calculating fold enrichment of positive hits (synergistic) or negative hits (antagonistic) over the total number of drugs in that category. Drug target analysis of the synergistic drug hits revealed significant enrichment in DNA synthesis and repair inhibitors, epigenetic modifiers, signal transduction and cytoskeletal inhibitors (Figure 2C). In contrast, drug target analysis of antagonistic drug hits revealed a higher enrichment in categories such as epigenetic modifiers, protein synthesis inhibitors, cytoskeletal inhibitors and proteasome inhibitors (Figure 2D).

**Figure 2.**
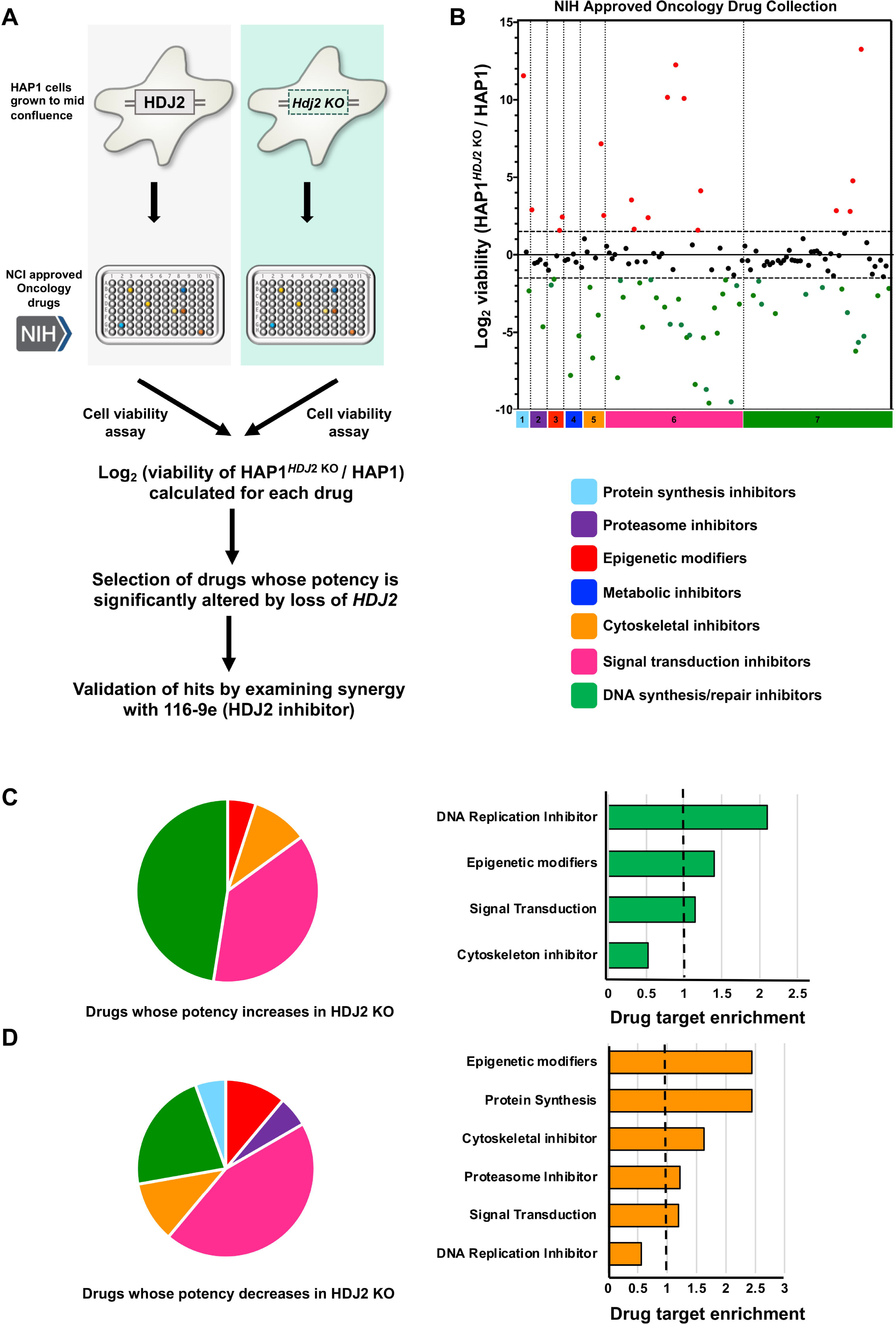
Sensitivity of WT and HDJ2 knockout cells to the NIH Approved Oncology Collection. (A) Workflow of high-throughput cell-based screen. (B) A collection of 132 drugs were screened at 50μmol/L with Wild-type and HDJ2 KO cells. Results are the average of at least triplicates and error is SEM. The dotted lines represent an interaction change of up or down two-fold. The dotted lines represent an interaction change of Log_2_ > 1.5 or Log_2_ < −1.5.The effect of drug combination are colored according to significant upregulation and downregulation: red (synergistic), green (antagonistic) or black (no significant change). C) & D) Drug ontology of synergistic and antagonistic hits based on the pathways affected by the approved oncology drugs in the screen.

Strikingly, compounds from different categories showed dissimilar distribution of log_2_ ratio of viability, implying that different pharmacologic mechanisms probably underlie the HDJ2 inhibitory capacity. Category no. 6 (Signal Transduction inhibitors) contained the most hits which were synergistic with HDJ2 loss. Loss of HDJ2 also substantially increased the potency of DNA synthesis and repair (DDR) inhibitors. These results are in agreement with our previous study showing that HDJ2 plays an important role in maintaining the stability of ribonucleotide reductase (RNR) complex which is important for DNA synthesis [13].

### Validation of anticancer drugs significantly altered for potency upon loss of HDJ2

Many anticancer compounds have low potency, poor therapeutic index or suffer from development of resistance [34]. Monotherapy is rarely efficient and instead drug cocktails are widely used in the clinic [23, 26]. Establishing these combinations can enhance the scope of preclinical studies and inform the design of future clinical trials. Although several compounds were identified as becoming significantly more potent in cells lacking HDJ2, it remained to be determined whether small molecule inhibition of HDJ2 could produce a similar result. Our previous bioinformatics analysis indicated that a large proportion of prostate cancer cells contain either amplification or mutation of HDJ2 (approximately 18%, see Figure 1). Therefore, we next analyzed the effect of treating prostate cancer cells (LNCaP) with a combination of 116-9e, a small molecule inhibitor of HDJ2 [35] and interesting hits from our screen. We decided to focus on three synergistic drugs discovered in the screen: cabozantinib (receptor tyrosine kinase inhibitor), clofarabine (an RNR inhibitor) [36] and vinblastine (microtubule inhibitor/G2 arresting agent) [37–40]. We also validated three drugs that demonstrated a significant loss of potency in cells lacking HDJ2: sorafenib (a VEGFR-2 inhibitor) [41], omacetaxine mepesuccinate (more commonly known as homoharringtonine, a protein translation inhibitor) [42] and idarubicin (topoisomerase II inhibitor) [43].To determine synergy in a quantitative manner, we calculated drug synergy (Combination Index values, CI) between 116-9e and either synergistic or antagonistic drugs hits across a broad range of concentrations using the Chou-Talalay method [44]. For three hits identified in our screen (cabozantinib, clofarabine and vinblastine) we confirmed significant synergy (CI<1) with 116-9e across a range of doses (Figure 3A, B & C). In contrast, idarubicin, omacetaxine and sorafenib displayed a significantly antagonistic interaction (CI>1) across a range of doses (Figure 3D, E & F).

**Figure 3.**
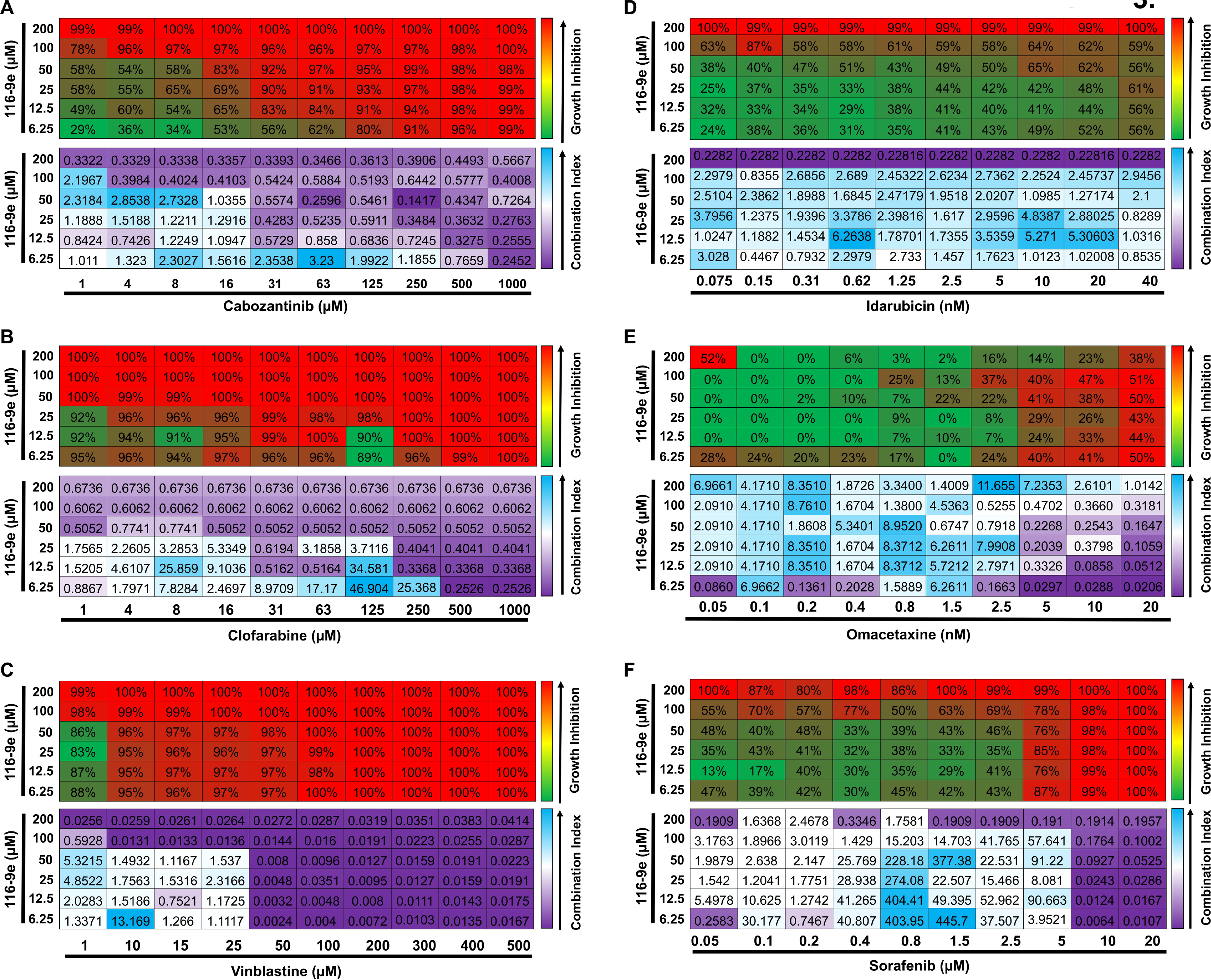
Drug interaction between 116-9e (HDJ2 inhibitor) and selected hits. LNCaP cells were treated with different concentration of Cabozantinib, Clofarabine, Vinblastine, Idarubicin, Omacetaxine and Sorafenib with or without 116-9e for 72 hours in RPMI-1640 medium containing 10% FBS. Each point is the mean ± SD for three independent experiments. Growth inhibition was determined using Cell Titer-Glo assay. Combination Index (CI, measure of drug synergy) was determined using Chou-Talalay method via Compusyn software. CI values of <1 indicate drug synergy.

These data suggest that while HDJ2 inhibition is a promising strategy to sensitize cells to some inhibitors, it might have inverse effects with other inhibitors.

### Evaluating the effects of dual targeting of identified drugs with HDJ2 inhibition on morphology and viability of prostate cancer spheroids

Recent studies have suggested that precision therapy approaches involving the exposure of drugs directly to the primary tumor tissue have the potential to augment the personalized medicine efforts and influence clinical decisions [45, 46]. Establishing *ex vivo* three-dimensional (3D) tumor spheroids or organoids derived from primary cancers can be easily established and potentially scaled to screen drug combinations [47]. These 3D cancer models appear to recapitulate features of the tumor of origin in terms of heterogeneity, cell differentiation, histoarchitecture, and clinical drug response and can be used for rapid drug screening [48]. We therefore next examined the effect of drug combination (three antagonistic and synergistic hits) on LNCaP spheroids. Specifically, changes in spheroid size and shape induced by the 3 antagonistic and synergistic drugs were determined. Visual examination revealed that for the synergistic drugs combination with 116-9e resulted in physical disruption of LNCaP spheroids, resulting in decrease in apparent spheroid size (Figure 4A). The disruption started on the second day of the treatment. However, when the 3 antagonistic drugs were administered along with 116-9e, there were minimal changes in spheroid morphology indicating that the combination was ineffective.

**Figure 4.**
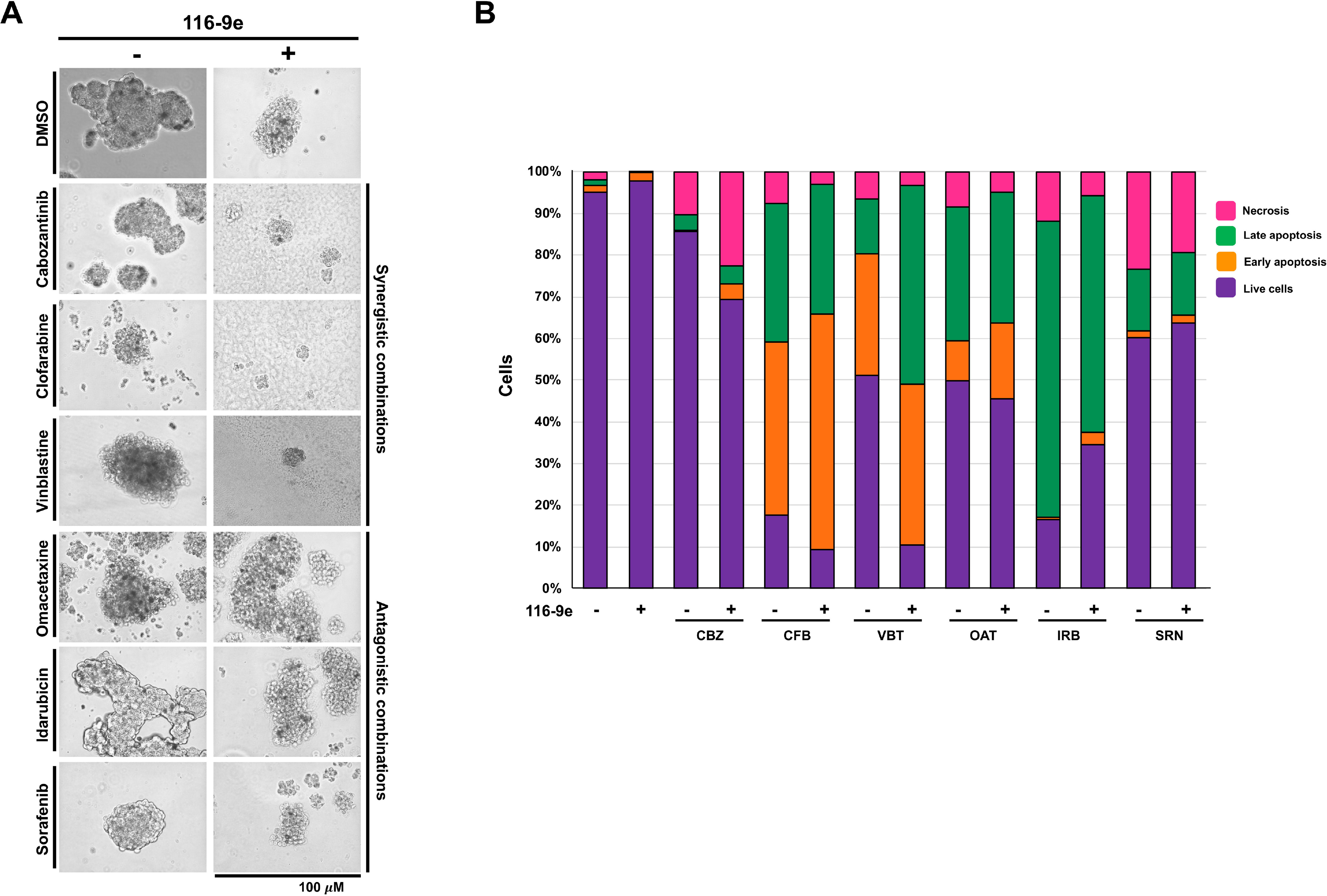
Effect of combination treatments on prostate cancer spheroids. A. Cells were plated on Matrigel coated 24 well plates. Six drugs (Cabozantinib, Clofarabine, Vinblastine, Idarubicin, Omacetaxine and Sorafenib) were tested in triplicates for prostate cancer spheroids. The pictures are representative images as acquired using EVOS cell imager. B. Proliferation of spheroids treated with Cabozantinib (CBZ), Clofarabine (CFB), Vinblastine (VBT), Idarubicin (IRB), Omacetaxine (OAT) and Sorafenib (SRN) measured using AnnexinV/PI staining.

Next, we measured the induction of apoptosis in the spheroids post drug treatments. We determined the kinetics of apoptosis induction using AnnexinV/PI staining. Drug-induced apoptosis was readily detected in the LNCaP spheroids treated with mono and dual drug combinations. In concurrence with the previous results, the combination of the three synergistic drugs with 116-9e displayed enhanced apoptosis as compared to the single drug treatment whereas spheroids treated with the 3 antagonistic drugs showed little or no difference in the rate of apoptosis as compared to the dual drug combination with 116-9e (Figure 4B).

## DISCUSSION

Although inhibitors of Hsp70 and Hsp90 have been developed for research purposes, conversion of these molecules for use in patient treatment have been hampered by toxicity issues [4]. We undertook this study to resolve conflicting literature on whether inhibiting HDJ2, a co-chaperone of Hsp70 may be useful as a novel anticancer strategy. Our bioinformatic analysis of HDJ2 expression and mutation clearly identify HDJ2 as being highly altered in a range of cancers, particularly in Prostate Cancer. This data in conjunction with a recent finding that Hsp40 is involved in functional regulation of ARv [22] makes HDJ2 inhibition an ideal choice as a novel therapeutic target in Prostate Cancer.

Chemogenomic screening of knockout cell lines produces both useful mechanistic and translational understanding of protein function. In this study, loss of HDJ2 increased the potency of a substantial number (31%) of clinically used anticancer drugs.

Hsp70 activates many proteins involved in the DNA damage response and DNA repair pathways (DDR). These include ATM, APE1, PARP1, XRCC1, LIG3, MSH2, MLH1 and Apollo [49–51]. In addition, studies from our group have established roles for both Hsp70 and HDJ2 in stability of the RNR complex [13, 50, 52]. As such, we would expect a high degree of synergy between loss of HDJ2 function and the DNA damage response/repair pathways. Correspondingly, around 20 commonly used anticancer DNA damage and Repair (DDR) inhibitors were found to be synergistic with loss of HDJ2. These included 5-fluorouracil (5-FU) and premetrexed, widely used anticancer drugs whose metabolites are incorporated into both DNA and RNA in addition to inhibiting thymidylate synthase [53]. Here we validated synergy with the RNR inhibitor clofarabine. Clofarabine is phosphorylated intracellularly to form cytotoxic active 5’-triphosphate metabolite, which inhibits the enzymatic activities of RNR and DNA polymerase, resulting in inhibition of DNA synthesis and repair[54]. In addition, we also identified PARP inhibitors olaparib and niraparib and the topoisomerase inhibitors etoposide, teniposide, valrubicin and dexrazoxane to have increased potency in our screen.

While most DDR inhibitors displayed increased potency with HDJ2 depletion, four of them were antagonistic to loss of HDJ2. These include topoisomerase inhibitors and nucleic acid synthesis inhibitors such as trifluridine, irinotecan, epirubicin (4’-epi-isomer of the antibiotic doxorubicin) and idarubicin (4-demethoxy analogue of daunorubicin)[55]. While at first these results seem paradoxical, it is worth noting that inrinotecan is a type I topoisomerase inhibitor, whereas Etoposide inhibits the type II class. It may be that Hsp70 and HDJ2 play different regulatory roles in the stabilization and activation of these related proteins. It should be noted that both idarubicin and epirubicin trigger TOPII-mediated DNA cleavage. The effects of these molecules may be prevented if HDJ2 alters the function of TOPII.

In addition to DDR, HDJ2 is also involved in signal transduction, with previous reports indicating that the yeast homologue of HDJ2 (Ydj1) is critical for supporting the integrity of kinase signaling networks [56]. HDJ2 is mobilized to specific sites within the nucleus in response to inappropriate targeting or folding of specific mutant receptors. HDJ2 overexpression ameliorates defective transactivation and trans repression activity of mutant Glucocorticoid receptors [57]. In line with the previous studies, we found that a handful of Receptor Tyrosine kinase inhibitors were synergistic with HDJ2 depletion. These included Vascular endothelial growth factor receptor (VEGFR) inhibitors such as sunitinib, cabozantinib, lenvatinib and pazopanib. Interestingly, randomized phase III clinical trials are being conducted to validate the efficacy of Cabozantinib in heavily pretreated prostate cancer patients [58]. One implication from our study is that HDJ2 inhibition might significantly enhance the effect of cabozantinib monotherapy.

Strikingly, some of the kinase inhibitors were antagonistic to HDJ2 depletion. These include VEGFR inhibitors such as regorafenib and sorafenib. This disparity can be explained by the different target receptors and mechanisms of action of these drugs. Interestingly, recent studies indicated that these small molecule inhibitors exhibit off-target effects. Some of these drugs are misidentified and mischaracterized for their target specific inhibition, which has contributed to the high failure rate of these drugs in treatment of cancer patients [59].

Other than its role in signal transduction, HDJ2 is also important for maintaining the cellular cytoskeleton. Previous studies have suggested that YDJ1 (yeast homolog of HDJ2) is important for the proper assembly of microtubules [60, 61]. Another report showed that HDJ2 depletion causes relocation of N-cadherin and enhanced activity of metalloproteinases. This leads to changes in the actin cytoskeleton indicating that HDJ2 is important for prevention of the amoeboid-like transition of tumor cells [62]. These studies indicated the involvement of HDJ2 in maintaining cytoskeletal organization. We found 3 anticancer drugs targeting the cytoskeleton to be synergistic with HDJ2 depletion, including vinblastine sulfate (cytoskeletal inhibitor that disrupts microtubule formation during mitosis and interferes with glutamic acid metabolism), estramustine (binds to microtubule-associated proteins (MAPs) and inhibits microtubule dynamics) and ixabepilone (promotes tubulin polymerization and microtubule stabilization, thereby arresting cells in the G2-M phase [63].

Strikingly, two of the tubulin inhibitors were found to be antagonistic to HDJ2 depletion. These include paclitaxel and ixabepilone. Paclitaxel inhibits the disassembly of microtubules resulting in the inhibition of cell division whereas Ixabepilone promotes tubulin polymerization and microtubule stabilization, arresting cells in the G2-M phase of the cell cycle [63]. This discrepancy again implies that these cytoskeletal inhibitors might have off target effects due to their mischaracterization [59].

Epigenetic modifying drugs display substantially modified potency depending on cellular HDJ2 status. While previous studies have indicated the association between proteomic changes and histone PTMs in response to Hsp90 inhibitor treatment in bladder carcinoma cells, no such association has been shown for HDJ2 and Histone PTMs [64]. Interestingly, vorinostat was the only drug that was synergistic to HDJ2 inhibition. It is a histone deacetylase inhibitor that binds to the catalytic domain of the histone deacetylases (HDACs) [65]. However, we identified two histone deacetylase inhibitor drugs to be antagonistic to HDJ2 depletion:panobinostat and romidepsin inhibit histone deacetylase (HDAC), inducing hyperacetylation of core histone proteins, which may result in modulation of cell cycle protein expression, cell cycle arrest in the G2/M phase and apoptosis [66]. This is the first study that indicates an association between histone PTMs and HDJ2. While these findings require further investigation, it is possible that HDJ2 regulates histone properties. Interestingly, both of the protein synthesis inhibitors (bortezomib and omacetaxine) in our screen were antagonistic to HDJ2 depletion [67]. We confirmed that omacetaxine (protein biosynthesis inhibitor) displayed reduced potency upon inhibition of HDJ2 [42].

Several important conclusions can be inferred from the data presented here. Firstly, our HTS screening method might be useful in the selection of drugs for individual patients in future studies, since the drug sensitivity of cancer cells is dependent on HDJ2 expression. For example, compounds that belong to the same category such as sunitinib and sorafenib may behave differently upon HDJ2 deletion.

Finally, in addition to intra-pathway synergistic combinations (VEGFRi, MAPKi pathway inhibitors, and DNA damage/cell-cycle checkpoint pathway combinations), which is consistent with a wealth of publications demonstrating intrapathway synergy [68, 69], we also discovered novel inter-pathway combinations of HDJ2. This study describes promising results and indicates an integrative approach based on HTS which has potential to govern cancer patient treatment by combination therapy. Taken together, this study suggests a potential Precision Medicine approach that has the potential to inform anticancer strategy based on patient HDJ2 status.

## Acknowledgements

The authors thank NIH for providing materials used in this study and J. Gestwicki, D. Dreau and T. Erick for helpful comments. This project was supported by NCI R15CA208773 and Grant-In-Aid of Research from Sigma Xi (G2018031591887158), The Scientific Research Society.

## Supplementary Data

Supplementary Table 1. Hits identified in combination screen with simultaneous treatment of Approved oncology drugs with HDJ2 Knockout HAP1 cell lines.

